# Global plant-frugivore trait matching is shaped by climate and biogeographic history

**DOI:** 10.1101/2021.12.17.473099

**Authors:** Ian R. McFadden, Susanne A. Fritz, Niklaus E. Zimmermann, Loïc Pellissier, W. Daniel Kissling, Joseph A. Tobias, Matthias Schleuning, Catherine H. Graham

**Affiliations:** Swiss Federal Institute for Forest, Snow and Landscape Research (WSL), Zürcherstrasse 111, 8903 Birmensdorf, Switzerland; Department of Environmental Systems Science, ETH Zürich, 8092 Zurich, Switzerland; Senckenberg Biodiversity and Climate Research Centre (SBiK-F), Senckenberganlage 25, 60325 Frankfurt am Main, Germany; Institut für Geowissenschaften, Goethe University Frankfurt, Altenhöferallee 1, 60438 Frankfurt am Main, Germany; Institute for Biodiversity and Ecosystem Dynamics (IBED), University of Amsterdam, P.O. Box 94240, 1090GE, Amsterdam, The Netherlands; Department of Life Sciences, Imperial College London, Silwood Park, Ascot SL5 7PY, UK

**Author notes:** Correspondence*: Ian R. McFadden, Swiss Federal Research Institute WSL, Zürcherstrasse 111, 8903 Birmensdorf, Switzerland. **Author emails:** SAF; NEZ; LP; WDK; JAT; MS; CHG. **Authorship statement:** Conceptualization: IRM, CHG, MS, JAT, WDK, SAF; data collection: JAT, MS, WDK, SAF; data analysis and visualization: IRM; writing (original draft): IRM; writing (reviewing and editing): IRM, CHG, JAT, MS, WDK, NEZ, SAF, LP.

**Keywords:** Functional biogeography, seed dispersal, birds, palms, structural equation modeling

## Abstract

Species interactions are influenced by the trait structure of local multi-trophic communities. However, it remains unclear whether mutualistic interactions in particular can drive trait patterns at the global scale, where climatic constraints and biogeographic processes gain importance. Here we evaluate global relationships between traits of frugivorous birds and palms (Arecaceae), and how these relationships are affected, directly or indirectly, by assemblage richness, climate and biogeographic history. We leverage a new and expanded gape size dataset for nearly all avian frugivores, and find a positive relationship between gape size and fruit size, i.e., trait matching, which is influenced indirectly by palm richness and climate. We also uncover a latitudinal gradient in trait matching strength, which increases towards the tropics and varies among zoogeographic realms. Taken together, our results suggest trophic interactions have consistent influences on trait structure, but that abiotic, biogeographic and richness effects also play important, though sometimes indirect, roles in shaping the functional biogeography of mutualisms.

## INTRODUCTION

Species interactions are important for structuring local communities and influence ecosystem function (Tilman 1982; Bertness & Callaway 1994; Callaway *et al*. 2002), but whether and how these interactions scale up to create global diversity patterns is currently debated. For example, Schemske et al. (2009) suggested that a gradient in the strength of species interactions is one cause of the latitudinal diversity gradient. This hypothesis has been challenging to test empirically because of a lack of data on species interactions, though some studies on this topic have used functional traits as proxies for interactions (Freeman *et al*. 2021). One way to overcome this challenge is to examine patterns of trait matching (Dehling *et al*. 2014), where pairs of traits of interacting taxa are compared to infer the strength of species interactions. However, understanding how species interactions are linked with trait variation at large scales is difficult using trait-based approaches alone, because phenotypic attributes of species involved in interactions may also be influenced by the abiotic environment (Burns 2004). Such environmental effects can act either directly on a species trait, or indirectly by altering the traits of the species with which it interacts (Maruyama *et al*. 2018). However, the extent to which trait matching is shaped by species interactions or by direct or indirect effects of climate, and how the strength of trait matching varies across the globe is not well understood.

Mutualistic interactions between fruiting plants and frugivores are a useful system to explore the effect of species interactions on macroecological patterns because seed dispersal can influence the structure and diversity of ecosystems at many scales (Levin *et al*. 2003; Bello *et al*. 2015; Gardner *et al*. 2019). Fruit and seed traits in plants and mandibular traits in frugivores are thought to be adaptations, at least in part, for seed dispersal (Janson 1983, Fig. 1, path g). Consequently, the functional relationship between plants and their dispersers is often reflected in a pattern of trait matching at local scales (Dehling *et al*. 2014). However, it is difficult to determine the underlying causes of trait matching at larger scales because the same traits may also be influenced by the abiotic environment. For example, the avian thermoregulation hypothesis (Tattersall *et al*. 2017) suggests that bird beak sizes will correlate positively with temperature as an adaptation for thermoregulation, via the heat shedding properties of large beaks (Fig. 1, path a). Conversely, the plant productivity hypothesis (Moles *et al*. 2007) states that more productive environments, such as tropical forests, will favor larger plants with larger seeds able to germinate in low light (Fig. 1, path d). Climatic constraints on traits involved in seed dispersal, such as mandible size in frugivores, can therefore act either directly on traits or indirectly by shaping the traits of the resources being consumed (e.g. Boag & Grant 1981).

**Figure 1.**
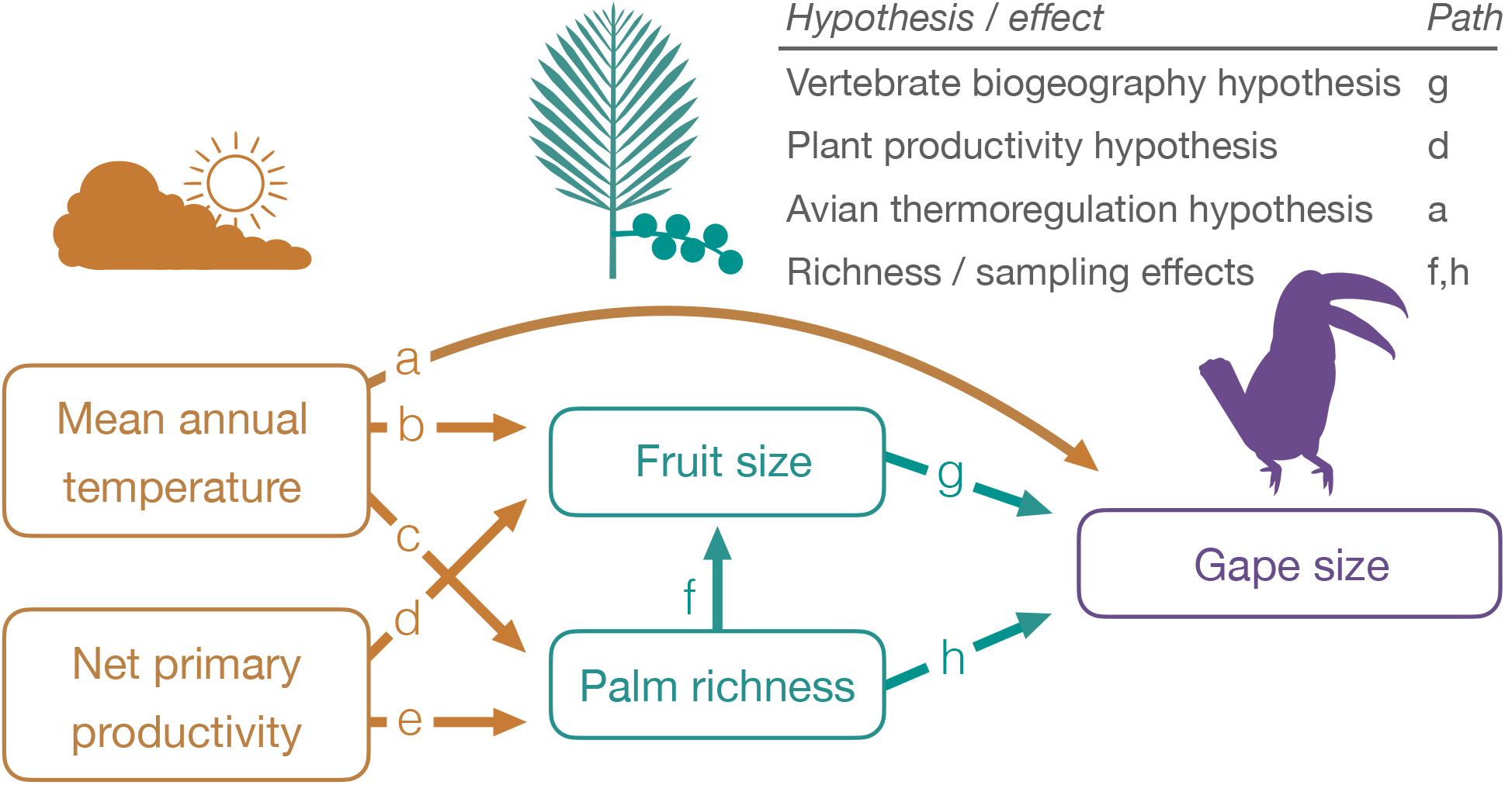
Hypothesized direct and indirect links between climatic variables, palm fruit size, palm richness and bird gape size, shown as a path diagram. Fruit size and palm richness can have direct effects on gape size (green arrows), while climate can have direct effects on fruit size and palm richness, as well as gape size, in addition to indirect effects on gape size through its effect on fruit size and palm richness (orange arrows). All relationships are predicted to be positive.

In addition to climatic constraints, global geographic variation in the size, foraging strategy and other functional attributes of vertebrate seed dispersers (Corlett & Primack 2011) may generate spatial variation in selective pressure on, and ecological sorting of fruiting plant species. This process may in turn lead to trait matching between fruits and frugivores among size-related, or other traits (Mack 1993; Kissling 2017; Sinnott-Armstrong *et al*. 2021). Testing this idea-hereafter the vertebrate biogeography hypothesis-requires quantifying traits, and trait matching across many biogeographic realms and a range of latitudes. Further, it may be expected that the largest observed trait values of animal and plant assemblages will be most correlated, as compared to averages, because frugivores cannot swallow fruits much larger than the width of the mandibles. This is particularly important for birds, which tend to swallow fruits whole (Wheelwright 1985).

Trait matching patterns may also be influenced by the richness of the plant and animal communities being compared (Maruyama *et al*. 2018, Fig. 1, paths f and h). For example, highly-diverse assemblages may be more likely to contain species with matching traits or extreme trait values due to sampling effects (Baraloto *et al*. 2010). In addition, species-rich assemblages also tend to be found in tropical forest regions, which have higher levels of dietary specialization among vertebrates (Belmaker *et al*. 2012) and often have long evolutionary histories (Davis *et al*. 2005) with ample time for diffuse coevolution, ecological sorting and ecological fitting (*sensu* Janzen 1985). This is important because mutualistic interactions such as seed dispersal occur across many regions that vary in richness (Sinnott-Armstrong *et al*. 2018), enabling analyses that infer the relative importance of biotic and richness effects on trait patterns.

To determine if a global signal of morphological matching exits and assess variability across zoogeographic realms in the functional biogeography of palm and bird traits, we integrate newly available trait datasets for birds and palms within a single multi-trophic framework (Fig. 1). As a test of the plant productivity hypothesis (Moles *et al*. 2007) and the avian thermoregulation hypothesis (Tattersall *et al*. 2017), respectively, we predict that fruit sizes will be larger in more productive regions (path d), and that regions with warmer temperatures will have birds with greater beak sizes (path a). Second, as a test of the vertebrate biogeography hypothesis (Mack 1993; Kissling 2017), we predict palm fruit size will correlate positively with bird gape size, and that these correlations will be stronger when comparing maximum trait values of assemblages as opposed to median values (path g). Finally, we predict assemblage richness will be positively related to gape and fruit size (paths f, h), and the range of gape and fruit sizes within assemblages, because diverse regions may be more likely to contain species with extreme trait values.

To test our predictions, we use palm fruit size and the gape sizes and beak volumes of frugivorous birds, which have previously been shown to be related at local and regional scales (Burns 2013; Chen & Moles 2015; Bender *et al*. 2018). To evaluate climatic influences on traits, we use mean annual temperature and net primary productivity (NPP). We focus here on palms because they are ubiquitous and ecologically important elements of many tropical and subtropical regions, and have likely interacted with frugivores for millions of years (Onstein *et al*. 2017, 2020). Similarly, birds are known to be important consumers of palm fruits (Benzing & Seemann 1989; Muñoz *et al*. 2019). We test our hypotheses using structural equation models as well as regression and residual analysis, and find that both climate and fruit size influence bird gape size, with climate having an indirect effect via its effect on fruit size and palm richness. In addition, we discover a positive relationship between gape size and fruit size globally, and a latitudinal gradient of increasing trait matching strength towards the equator.

## METHODS

### Functional trait data

To test for plant-frugivore trait matching at the global scale, we used bird gape size and beak volume, and palm fruit size. To assemble trait measurements for frugivorous birds, we compiled a dataset of beak measurements taken from wild-caught and released individuals, as well as specimens accessed in numerous museums and research collections worldwide (Tobias *et al*. 2021). We defined frugivores as those species with a diet containing 50% or more fruit, and only kept species matching this criterion. We attempted to measure at least two males and two females of each frugivore species, with the final dataset containing measurements from 12925 individuals and a total of 1129 species. For beak volume and diet traits, we used data from Pigot et al. (2020) and Tobias et al. (2021). To quantify beak volume (*n* = 1129 spp.), we multiplied beak length, measured from the tip of the beak along the culmen to the base of the skull, with beak width and beak depth, both measured at the anterior edge of the nostrils (units = mm3).

To quantify avian frugivore gape size (previously termed ‘gape width’ or ‘beak width’ in some studies), we used data from Pigot *et al*. (2016), Bender et al. (2018) and Hanz et al. (2019), and also collected previously unpublished measurements on thousands of additional individuals. We defined gape size as the horizontal width of the beak measured between the points at which the upper and lower mandibles meet. This trait is important because birds most often swallow fruits whole, and thus gape size should tend to set an upper bound to fruit sizes birds can consume (Wheelwright 1985; Burns 2013). Unlike standard beak measurements, such as those in Tobias et al. (2021), the global dataset of gape size is unique to this study (doi.org/10.5061/dryad.tqjq2bw05). We include gape size data in two ways. First, as species averages used in our models, and second, by providing the underlying data from all museum specimens and wild-caught individuals along with metadata on source, collection locality and measurer. We found that gape size and beak volume are strongly correlated at the botanical country scale (Fig. S1A, *r* = 0.95), so we chose to include only gape size in our main analyses because of its strong link with maximum ingestible fruit size.

For palm fruit size, we used the PalmTraits database (Kissling *et al*. 2019), which contains vegetative and reproductive traits for nearly all palm species. We first removed three species not thought to be dispersed by animals (Dransfield *et al*. 2008): the coconut (*Cocos nucifera*), the coco de mer (*Lodoicea maldivica*) and the nipa palm (*Nypa fruticans*). Then, we extracted species mean fruit width (*n* = 1992 spp.) and length (*n* = 2049 spp.), which were highly correlated (*r* = 0.87). Next, we combined fruit width and length measurements to obtain a measure of fruit size (*n* = 2051 spp.). Fruit width was used when available, and for species with only length data (*n* = 59) we estimated fruit width via the allometric relationship between length and width for species in PalmTraits having both traits, using the equation log(fruit width) = 1.026 × log(fruit length) – 0.337. Fruit size, rather than seed size, was used because this trait should tend to limit the gape size of birds foraging on fruits, especially because most palms have only one seed (Dransfield *et al*. 2008).

### Species distribution data

To determine how biogeographic history, species richness and climate influence global-scale trait matching, we compiled geographic distribution data for birds and palms, and aggregated these data via several methods into the same geographic units (Fig. S2). Geographic range maps of birds were extracted from the BirdLife International database (Birdlife International and NatureServe 2013) and palm distribution data from the PalmTraits database (Kissling *et al*. 2019). As with other plant clades, detailed range maps for many palm species are not available, thus species distributions in the PalmTraits database are specified as presences or absences within botanical countries, which are standardized regions defined by the International Taxonomic Databases Working Group (TDWG, Brummitt *et al*. 2001). To match the scale of palm distribution data, assemblages of bird species were defined by overlapping the bird ranges with the botanical country polygons, thus creating species lists for each botanical country. We removed the Rufous-necked hornbill (*Aceros nipalensis*) from Tibet, as this species is known to be a vagrant in this region. We then calculated the maximum and median trait values for each botanical country using all species occurring within the botanical country. We excluded from further analysis regions with less than three species, because trait matching can only be quantified in interacting assemblages of several species. Finally, we assigned botanical countries to zoogeographic realms (*sensu* Holt *et al*. 2013), placing islands not previously classified this way within the realm most geographically close to the island. Figures S3-S5 show the variation across botanical countries in both single size trait values and richness for bird and palms.

### Climatic data

We obtained climatic data from several sources, including NPP derived from MODIS (Running *et al*. 2004) averaged over the years 2001-2011, and mean annual temperature from CHELSA (Karger *et al*. 2017). Though NPP and MAT are somewhat interrelated, we retained both variables because each is related to a climatic hypothesis we test, i.e., the plant productivity and avian thermoregulation hypotheses, respectively, and because the correlation between them is weak in our dataset (Fig. S1). Climatic data was then aggregated in the same way as trait data for botanical countries: maximum and median MAT and NPP values were calculated for each botanical country containing bird and palm species included in our analysis.

### Structural equation modeling

To determine how direct and indirect effects can create or obscure trait matching, and how these effects vary across zoogeographic realms, we used piecewise structural equation models (piecewise SEMs, Lefcheck 2016). Piecewise SEMs differ from classic path analyses because each component model is solved separately instead of using a single variance-covariance matrix, which allows for greater flexibility in the specification of each path relationship. After log-transforming and scaling each continuous variable (to μ = 0, σ = 1), we used linear mixed effects models to fit model paths between plant traits, bird traits and climate, specifying in each case zoogeographic realm as a random effect. We used the maximum trait and climatic values of each botanical county in the main model, however we also fit the model with median values to compare results using the centers of distributions. We then extracted standardized model coefficients for each direct and indirect path, as well as the marginal *R*^2^-the variance explained by fixed factors, and the conditional *R*^2^-the variance explained by both fixed and random, here zoogeographic realm, effects.

### Trait mapping and regression analysis

To visualize trait matching globally, we co-plotted the maximum values of gape and fruit size for each botanical country in space and via linear regressions. We also incorporated sampling effects due to species richness by fitting linear regressions of size traits with richness for both birds (log gape size ~ log bird richness) and palms (log fruit size ~ palm richness). We then used the residuals of these regressions to plot a richness-corrected relationship between gape and fruit size. Next, we used model II regression to test for significant deviations in these relationships from the 1:1 line, i.e., the line of isometry. Finally, to test whether diverse regions are more likely to contain species with extreme trait values, which may influence our path model via palm richness (Fig. 1, paths f-h), we quantified for each botanical country the range of trait values (max - min) for bird gape size and palm fruit size. We then fit linear and 2^nd^ degree polynomial regressions of fruit size range and gape size range with palm richness, and between bird gape size range and palm fruit size range across all botanical countries.

To understand how trait matching varies across latitude and among realms, we extracted the residuals of the linear regression between fruit and gape size, as well as fitted values of gape size. We interpret small residuals for a given botanical country as an indication of strong trait matching. Compared to regions where residuals are large, i.e., traits are more mismatched, small residuals suggest fruit size is predictive of, or matched to, gape size in this region. We then examined the relationship between residuals, i.e., trait matching, and latitude, compared average residual values among zoogeographic realms and plotted residuals in space for all botanical countries.

## RESULTS

### Structural equation models

Using a structural equation modeling approach, we found that climate, acting indirectly via effects on palm richness, is an important predictor of bird gape size (Fig. 2, see Fig. S6 for SEM using median trait values). For example, temperature was significantly and positively linked to both fruit size and richness (Fig. 2). However, we did not find support for the avian thermoregulation hypothesis, as temperature was negatively related to gape size (Fig. 2), and to beak volume when used instead of gape size in the SEM (not shown, see Fig. S1A for pairwise correlations). Counter to the plant productivity hypothesis, NPP was not significantly related to fruit size, though it was linked indirectly via an effect on palm richness (Fig. 2). In addition to indirect effects, there was a direct effect of fruit size on gape size (Fig. 2), both in models of maximum and median fruit and gape sizes (Fig. 2, Fig. S6). Our results also indicate large spatial variation in trait matching across zoogeographic realms: conditional *R^2^* values incorporating realm as a random effect were substantially higher, on average explaining 27% more variance than marginal *R*^2^ values which include only fixed effects. When we added botanical countries with 1-2 species to the analysis, the link between fruit size and gape size became non-significant (Fig. S7A), which may reflect limited trait-matching in assemblages containing very few species.

**Figure 2.**
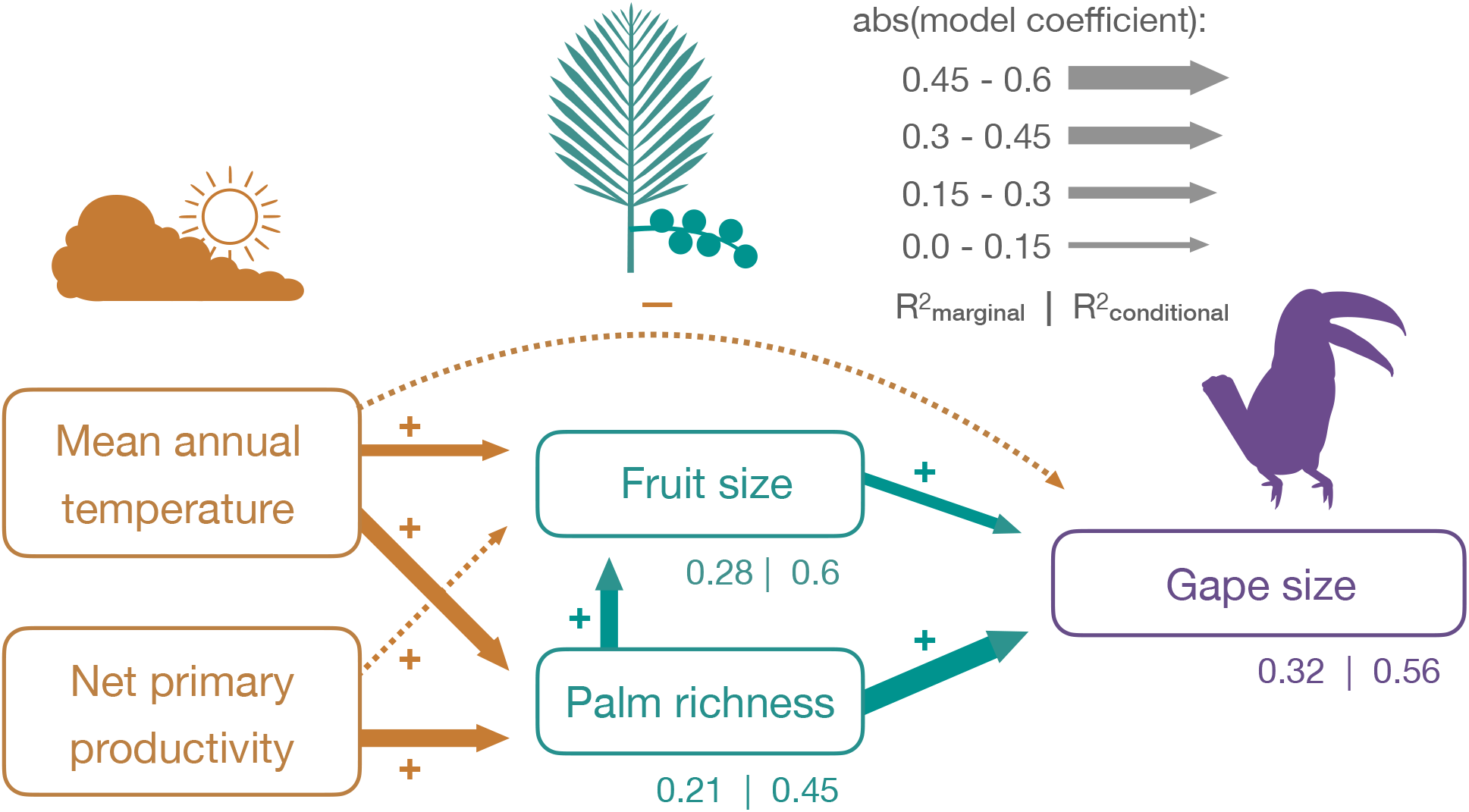
Path diagram showing how climate, fruit size and palm richness affect bird gape size. For all variables the maximum observed value of each botanical country (see *Methods*) was used, and Wallace realm (Holt *et al*. 2013) was specified as a random effect in the piecewise SEM sub-models. Line widths are scaled to the absolute value of model coefficients, solid lines indicate significant and dotted lines non-significant relationships. See Table S1 for path coefficients and Fig. S6 for a path model fit with median values for botanical countries.

### Trait mapping and regression analysis

Gape and fruit size trait values broadly co-varied in space and exhibited a high degree of spatial clustering (Fig. 3A). For example, gape and fruit sizes were both lowest in the Southeast United States and highest in Southeast Asia, while tropical South America and much of Africa tended to have intermediate values of both traits. In contrast, islands tended to have trait values that differed to a large degree from one another, i.e., were less matched. Overall, we found a positive relationship between maximum fruit and gape sizes across botanical countries (Fig. 3B, *r* = 0.57, *p* < 0.001), and similarly for residuals from regressions of bird and palm traits with assemblage richness (Fig. 3C, *r* = 0.34, *p* < 0.001). For both relationships (Fig. 3B&C), model II regression analysis demonstrated that the observed slope was flatter than the 1:1 line (non-residual values: slope = 0.62, 95% CI: 0.54-0.71; residual values: slope = 0.42, 95% CI: 0.35-0.49). Thus, best fit lines fell closer to the fruit size axis than the gape size axis in both cases, suggesting that the largest palm fruits were on average bigger than the widest bird beaks. As predicted, when median trait values of botanical countries were used the relationship between gape and fruit size was somewhat weaker (*r* = 0.28, *p* = 0.001, Fig. S1B). Results were similar when species poor botanical countries with 1-2 species were included in the analysis (Fig. S7B&C). Lastly, we found the range of fruit size and gape size traits within botanical countries increased with assemblage richness (Fig. 4A&B, *R^2^* = 0.43 and 0.25, respectively; all *p* < 0.001) and that the ranges of both traits were positively related to one another (Fig. 4C, *R*^2^ = 0.31, *p* < 0.001). Botanical countries with low palm richness had small to large gape size ranges, while areas with high palm richness had only large gape size ranges (Fig. 4B).

**Figure 3.**
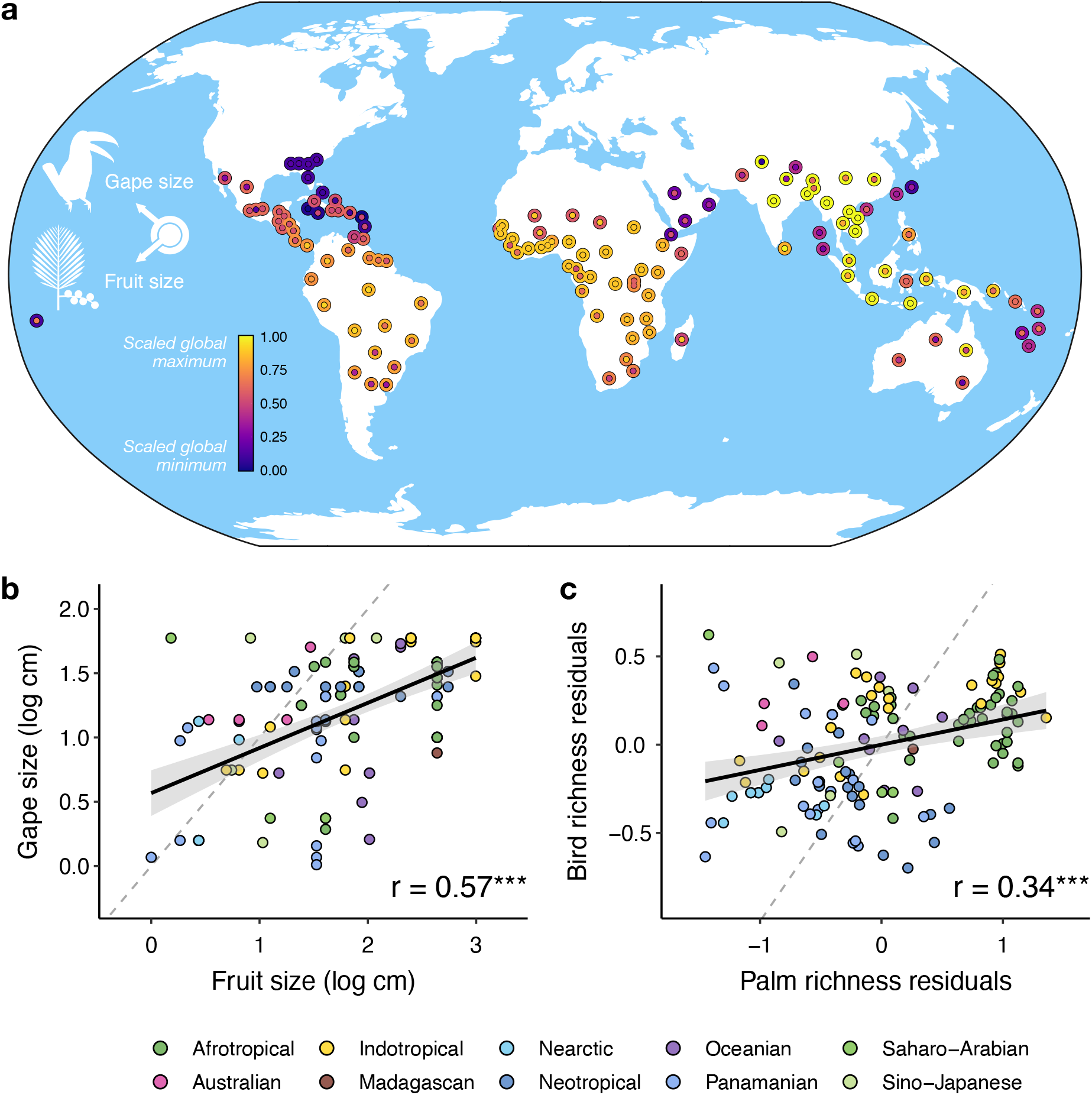
Global associations between palm fruit size and bird gape size. (a) Spatial variation in bird and palm trait matching across botanical countries (see *Methods*). Outer ring of each point is colored by gape size while inner points are colored by fruit size, warmer colors indicate higher values. Values are the maximum for each botanical country, logged and rescaled to 0-1, plotted at the centroid position. See Fig. S2 for names of each botanical country. (b) Fruit and gape size are correlated at the global scale, as are residuals calculated from a regression with log-transformed gape and fruit size and log transformed bird and palm richness values for each botanical country (c, see *Methods*). Points are colored according to Wallace realm (*sensu* Holt *et al*. 2013), dashed line is line of isometry, *** = p < 0.001.

**Figure 4.**
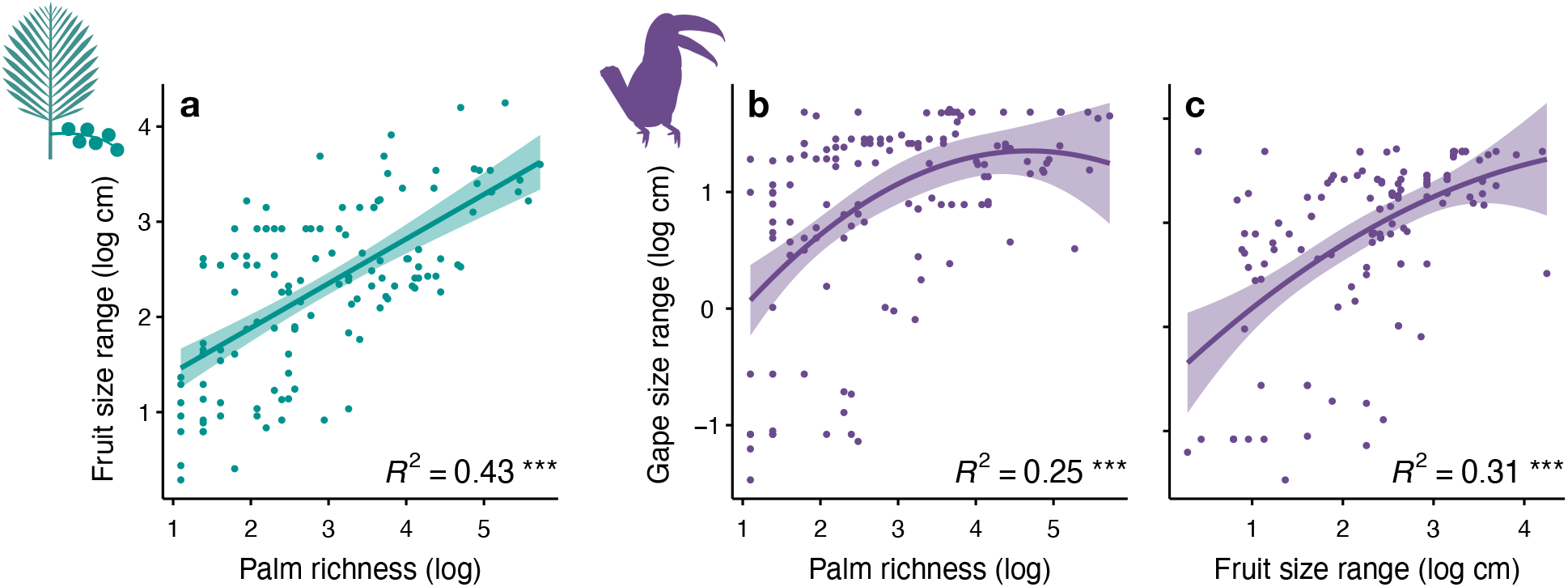
Effect of assemblage richness on trait ranges. (a) The range of fruit sizes within botanical countries (see *Methods*) increases with palm richness. (b-c) The range of gape sizes within botanical countries also increases with both palm richness and fruit size range. Lines and 95% confidence intervals are from linear (a) and 2^nd^ degree polynomial (b-c) fits. *** = p < 0.001.

Trait matching, as measured via residuals from the relationship between gape size and fruit size, significantly decreased away from the equator, i.e., residuals were larger further away from the equator (Fig. 5A). The major tropical forest-dominated realms-Afrotropical, Indotropical, and Neotropical-all tended to have comparatively strong trait matching, i.e., small average residual values (mean = 0.11, Fig. 5B). However, these regions differed in whether fruit size or gape size was larger than expected (negative vs. positive average residual values, respectively). Island and desert regions tended to be more mismatched in their traits (e.g., Madagascar, Oceania, the Arabian Peninsula) than most other realms, often due to larger fruit sizes compared to gape sizes (Fig. 5C). We found similar results when botanical countries with 1-2 species were included in the analysis (Fig. S7D&E).

**Figure 5.**
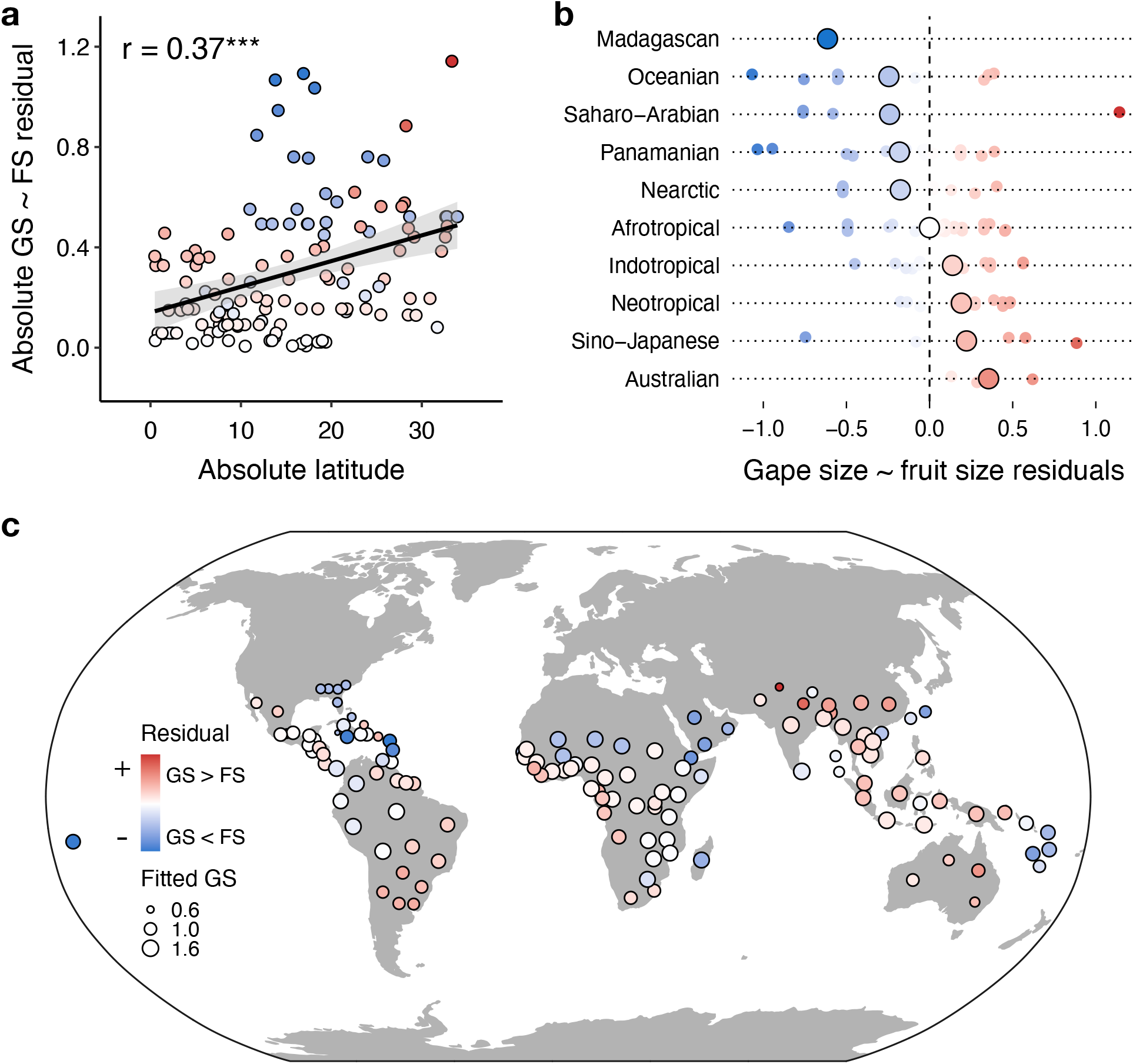
Global matching patterns of gape size (GS) and fruit size (FS). (a) Absolute residual values from a regression between log-transformed gape and fruit size (see Fig. 3B) decrease towards the equator (absolute latitude = 0), suggesting the strength of trait matching is strongest in equatorial regions. More red colors indicate a location has frugivores with larger gape sizes than predicted by fruit size, and more blue colors the opposite. (b) Trait matching strength differs among zoogeographic realms, as quantified via residuals from the gape size ~ fruit size regression (see Fig. 3B). (c) Global spatial variation in the residuals of the gape size ~ fruit size regression used to make panels a and b, with points colored as in panels a and b, and point size scaled to the fitted value of gape size from the same regression (Fig. 3B). See Fig. S2 for names of each botanical country.

## DISCUSSION

In our global analysis of bird and palm trait matching, we find i) a positive association between fruit size and gape size, with significant variation among zoogeographic realms, ii) that indirect effects of climate on trait matching are more important than direct effects, and iii) a latitudinal gradient in the strength of trait matching, increasing towards the tropics. Our study provides evidence that trait matching is likely created not only by traits of interacting partners, but also by indirect climatic influences on species phenotype. In addition, assemblage richness was an important factor shaping trait patterns, suggesting that sampling effects caused by geographic variation in richness, or unmeasured variation that covaries with richness, can influence large-scale trait matching patterns. In total, our results suggest that trait matching consistently emerges from seed dispersal interactions over large areas, but that sampling effects, climatic variation and biogeographic history have important, but often indirect, contributions to the functional biogeography of mutualisms.

It is debated whether biotic interactions influence plant and animal traits and assemblage structure in a consistent way at the global scale (Sinnott-Armstrong *et al*. 2018; Dugger *et al*. 2019). Using a new and expanded dataset of gape sizes for nearly all frugivorous bird species, we found a positive relationship between bird gape and fruit size globally, suggesting seed dispersal interactions are generally important drivers of trait patterns and thus of the ecosystem functions these interactions mediate (Harrison *et al*. 2013; Bello *et al*. 2015). Nevertheless, historical and environmental factors were also important. Our results add to a growing body of evidence that biotic interactions play an important role in generating broad-scale patterns of trait distributions (Maruyama *et al*. 2018; Hargreaves *et al*. 2019). For example, a positive relationship between mammal body mass and palm fruit size was also recently found (Lim et al. 2020), suggesting seed dispersal mutualisms are generally important in shaping trait patterns of both plants and frugivores.

Though we show species traits and climatic factors are important drivers of trait matching, the form and strength of this relationship may be influenced by the number and type of interacting species (Šímová *et al*. 2013) and by unmeasured interactions. For example, in our SEM the effect of palm richness on gape size was significant for assemblages of three or more species. However, this relationship was not significant when including botanical countries with only 1 or 2 species (Fig. S7). This result suggests that trait matching is less likely to occur if there are too few species with which to match. Also, interactions between birds and fruits are known to be diffuse in seed dispersal networks, and the diffuse nature of these interactions may make 1:1 trait matching unlikely (Guimarães *et al*. 2017). Indeed, our results indicated maximum sizes of palm fruits tended to be larger than maximum bird gape sizes. One explanation for this result is that palms are also dispersed by mammalian frugivores and omnivores, which tend to have larger body sizes than birds (French & Smith 2005), especially when including extinct megafauna (Lim *et al*. 2020).

We predicted that there is a higher chance of sampling extreme trait values in species-rich assemblages, which may increase the probability of trait matching through evolutionary dynamics (Onstein *et al*. 2017), ecological fitting (Janzen 1985) and biotic specialization (Maruyama *et al*. 2018). In line with the prediction, we found that as botanical countries became more species rich the range of trait values within them increased, suggesting that sampling effects increase the strength of trait matching. Therefore, trait matching appears more likely to emerge in species rich assemblages, which reinforces our finding that trait matching strength increases towards the equator (Fig. 5A), where biodiversity often peaks. One potential mechanism for these patterns is that character displacement and specialization on specific partners is more likely to occur in species rich areas (Maruyama *et al*. 2018), which also may have high functional diversity (Dehling *et al*. 2014). Future studies should account for variability in species richness in large-scale analyses of trait matching and explore in more depth the underlying mechanism of how richness shapes trait matching patterns.

Many traits involved in mutualisms are thought to be also influenced by abiotic factors (Burns 2004; Sales *et al*. 2021). However, using our SEM framework we found little support for direct climatic constraints on fruit and beak traits, as suggested by the plant productivity (Moles *et al*. 2007) and avian thermoregulation hypotheses (Tattersall *et al*. 2017). Thus, biotic effects such as traits involved in seed dispersal mutualisms (e.g. Galetti *et al*. 2013) and sampling effects of assemblage richness may be more important than thermoregulation or ecosystem productivity in determining global-scale trait patterns. Nonetheless, we only focus here on palms and frugivorous birds, and cannot make inference about the validity of the abiotic hypotheses for other clades and functional groups. However, our results do provide evidence for indirect effects of climate on plant and animal traits, and thus both direct and indirect effects should be assessed in future studies of trait biogeography, particularly for traits thought to shape mutualistic interactions.

A latitudinal gradient in the strength of species interactions has long been suggested (Schemske *et al*. 2009), though tests have been scarce (Hargreaves *et al*. 2019; Freeman *et al*. 2021). Here, we find that the strength of trait matching in the seed dispersal mutualism increases towards the equator (Fig. 5A), in line with previous results which found increased morphological matching towards low latitudes in pollination mutualisms (Sonne *et al*. 2020). Our results may reflect higher levels of trophic specialization in tropical ecosystems (Belmaker et al. 2012) or the greater evolutionary age of tropical clades, and thus the longer period of time available for diffuse trait coevolution to generate matching traits among interacting partners (Davis *et al*. 2005; Schemske *et al*. 2009). As mentioned above, trait matching may also be more likely to occur in diverse tropical assemblages with many interacting species because some species may possess matching traits by chance, or because trait matching emerges more rapidly due to the high number of interactions.

In contrast to latitudinal patterns, we found that the strength of trait matching in seed dispersal mutualisms was less consistent among zoogeographic regions. For example, we find Southeast Asia has some of the largest frugivore gape sizes globally, driven largely by hornbills (Bucerotidae). In contrast, mammal body sizes of extant and extinct species in Southeast Asia tend to be small compared to other regions such as Africa (Lim *et al*. 2020). The degree of matching outside tropical regions also varies idiosyncratically. For example, we found greater mismatching in some island and other depauperate areas compared to the equatorial tropics. This result could be due to the effects of isolation on islands filtering out taxa that cannot disperse to them (Yap *et al*. 2018), or the environmental filtering of non-adapted taxa from deserts or other xeric areas, or both.

Some important factors may affect our results due to the complex nature and global scale of the datasets used. First, the spatial resolution of palm distribution data required aggregating bird ranges and climatic data to the same scale, that of botanical countries. Therefore, the spatial units in our analysis were larger than traditionally defined communities of interacting species, here representing regional assemblages. Second, it should be noted that trait matching within a region does not imply co-occurring species are necessarily interacting partners (Blanchet *et al*. 2020). In our analysis, we could not include interaction data due to a lack of sufficient coverage over large areas (Poisot *et al*. 2021). Third, while birds are ecologically important dispersers of palms, several other groups also feed on palms, including mammals, lizards and insects (Benzing & Seemann 1989). Conversely, the diets of frugivorous birds often include other fruit species in addition to palms, and may also include insects and small mammals (Corlett & Primack 2011). Despite these potential constraints, we find predictable structure in trait patterns both in terms of trait matching across trophic levels and a consistent latitudinal gradient in the strength of this relationship.

### Conclusions

In summary, we find evidence for gape and fruit size trait matching between birds and palms at the global scale, suggesting that the seed dispersal mutualism influences and constrains the size of both plant and frugivore traits. We also find that climate can influence bird traits via links with plant richness, though not directly as has been hypothesized (Moles *et al*. 2007; Tattersall *et al*. 2017). These results contribute to our understanding of how biotic interactions influence diversity at large scales through both direct and indirect effects, with large-scale analyses across trophic levels now increasingly possible thanks to open, global trait data (Kissling *et al*. 2019; Kattge *et al*. 2020; Tobias *et al*. 2021). This study, and future research on other clades, could be used to predict how biotic interactions mediate global-change impacts on biodiversity (Schleuning *et al*. 2020) and to what extent these interactions are themselves vulnerable to human impacts from biotic (e.g., introduced species, species loss) and abiotic factors such as climate change (Tobias *et al*. 2020). Finally, the analyses presented here should be expanded to include all clades involved in seed dispersal mutualisms, supported by the continued collection of large plant and animal trait databases, to gain a more holistic understanding of the role trophic interactions play in generating ecosystem function and resilience.

## ACKNOWLEDGMENTS

We are grateful to Adam Devenish and Julie Marin for help with data organization, and also to all measurers of wild-caught birds, bird specimens and the respective collection holders who have been key to assembling the global bird trait dataset (Tobias et al. 2021) and the gape size dataset provided with this paper (doi.org/10.5061/dryad.tqjq2bw05). We thank members of the WDK, CHG, MS and SAF research groups for constructive feedback on the work, and Jeremy Kerr and two anonymous reviewers for helpful comments. Bird trait data collection was supported by the UK Natural Environment Research Council (Nos. NE/I028068/1 and NE/P004512/1 to JAT) and by the German Research Foundation (Nos. SCHL 1934/3-1 and SCHL 1934/2-3 to MS). IM, CG were supported by the European Research Council (ERC) under the European Union Horizon 2020 research and innovation program (No. 787638 granted to CG) and the Swiss National Science Foundation (No. 173342 granted to CG). WDK acknowledges funding from the Netherlands Organization for Scientific Research (No. 824.15.007) and University of Amsterdam (via a starting grant and Faculty Research Cluster ‘Global Ecology’). SAF was funded by the German Research Foundation (FR 3246/2-2) and the Leibniz Association (Leibniz Competition P52/2017). This publication is a contribution of the Swiss WSL Institute Biodiversity Center.

## Data accessibility statement

Bird beak and gape size trait data at the species and individual level, with associated metadata, is available via Dryad (doi.org/10.5061/dryad.tqjq2bw05). Aggregated data on bird and palm traits as well as climate at the level of botanical countries, with code to reproduce the results, is accessible via Zenodo repository (doi.org/10.5281/zenodo.5494437). Palm trait data at the species level is from PalmTraits (doi.org/10.5061/dryad.ts45225) and global climatic data is from CHELSA (chelsa-climate.org).

## REFERENCES

Baraloto, C., T. Paine, C.E., Patino, S., Bonal, D., Hérault, B. & Chave, J. (2010). Functional trait variation and sampling strategies in species-rich plant communities. Funct. Ecol., 24, 208–216.

Bello, C., Galetti, M., Pizo, M.A., Magnago, L.F.S., Rocha, M.F., Lima, R.A.F., et al. (2015). Defaunation affects carbon storage in tropical forests. Sci. Adv., 1, e1501105.

Belmaker, J., Sekercioglu, C.H. & Jetz, W. (2012). Global patterns of specialization and coexistence in bird assemblages. J. Biogeogr., 39, 193–203.

Bender, I.M.A., Kissling, W.D., Blendinger, P.G., Böhning-Gaese, K., Hensen, I., Kühn, I., et al. (2018). Morphological trait matching shapes plant-frugivore networks across the Andes. Ecography, 41, 1910–1919.

Benzing, D.H. & Seemann, J. (1989). A review of animal-mediated seed dispersal of palms. Selbyana, 2, 133–148.

Bertness, M.D. & Callaway, R. (1994). Positive interactions in communities. Trends Ecol. Evol., 9, 191–193.

Birdlife International and NatureServe. (2013). Bird species distribution maps of the world. BirdLife International, Cambridge, UK.

Blanchet, F.G., Cazelles, K. & Gravel, D. (2020). Co-occurrence is not evidence of ecological interactions. Ecol. Lett., 23, 1050–1063.

Boag, P.T. & Grant, P.R. (1981). Intense natural selection in a population of Darwin’s finches (geospizinae) in the Galápagos. Science, 214, 82–85.

Brummitt, R.K. (2001). World Geographical Scheme for Recording Plant Distributions Edition 2. Hunt Institute for Botanical Documentation, Pittsburgh; 2001. Group. International working group on taxonomic databases for plant sciences (TDWG), Pittsburgh.

Burns, K.C. (2004). Scale and macroecological patterns in seed dispersal mutualisms. Glob. Ecol. Biogeogr., 13, 289–293.

Burns, K.C. (2013). What causes size coupling in fruit-frugivore interaction webs? Ecology, 94, 295–300.

Callaway, R.M., Brooker, R.W., Choler, P., Kikvidze, Z., Lortie, C.J., Michalet, R., et al. (2002). Positive interactions among alpine plants increase with stress. Nature, 417, 844–848.

Chen, S.C. & Moles, A.T. (2015). A mammoth mouthful? A test of the idea that larger animals ingest larger seeds. Glob. Ecol. Biogeogr., 24, 1269–1280.

Corlett, R.T. & Primack, R.B. (2011). Tropical Rain Forests: An Ecological and Biogeographical Comparison: Second Edition. John Wiley & Sons.

Davis, C.C., Webb, C.O., Wurdack, K.J., Jaramillo, C.A. & Donoghue, M.J. (2005). Explosive radiation of Malpighiales supports a mid-cretaceous origin of modern tropical rain forests. Am. Nat., 165, E36–E65.

Dehling, D.M., Töpfer, T., Schaefer, H.M., Jordano, P., Böhning-Gaese, K. & Schleuning, M. (2014). Functional relationships beyond species richness patterns: Trait matching in plant-bird mutualisms across scales. Glob. Ecol. Biogeogr., 23, 1085–1093.

Dransfield, J., Uhl, N.W., Asmussen, C.B., Baker, W., Harley, M.M. & Lewis, C.E. (2008). Genera Palmarum : the evolution and classification of palms. Kew Publishing.

Dugger, P.J., Blendinger, P.G., Böhning-Gaese, K., Chama, L., Correia, M., Dehling, D.M., et al. (2019). Seed-dispersal networks are more specialized in the Neotropics than in the Afrotropics. Glob. Ecol. Biogeogr., 28, 248–261.

Freeman, B.G., Weeks, T., Schluter, D. & Tobias, J.A. (2021). The latitudinal gradient in rates of evolution for bird beaks, a species interaction trait. Ecol. Lett. (this volume).

French, A.R. & Smith, T.B. (2005). Importance of body size in determining dominance hierarchies among diverse tropical frugivores. Biotropica J. Biol. Conserv., 37, 96–101.

Galetti, M., Guevara, R., Côrtes, M.C., Fadini, R., Von Matter, S., Leite, A.B., et al. (2013). Functional extinction of birds drives rapid evolutionary changes in seed size. Science, 340, 1086–1090.

Gardner, C.J., Bicknell, J.E., Baldwin-Cantello, W., Struebig, M.J. & Davies, Z.G. (2019). Quantifying the impacts of defaunation on natural forest regeneration in a global meta-analysis. Nat. Commun., 10, 1–7.

Guimarães, P.R., Pires, M.M., Jordano, P., Bascompte, J. & Thompson, J.N. (2017). Indirect effects drive coevolution in mutualistic networks. Nature, 550, 511–514.

Hargreaves, A.L., Suárez, E., Mehltreter, K., Myers-Smith, I., Vanderplank, S.E., Slinn, H.L., et al. (2019). Seed predation increases from the Arctic to the Equator and from high to low elevations. Sci. Adv., 5, eaau4403.

Harrison, R.D., Tan, S., Plotkin, J.B., Slik, F., Detto, M., Brenes, T., et al. (2013). Consequences of defaunation for a tropical tree community. Ecol. Lett., 16, 687–694.

Holt, B.G., Lessard, J.-P., Borregaard, M.K., Fritz, S.A., Araújo, M.B., Dimitrov, D., et al. (2013). An update of Wallace’s zoogeographic regions of the world. Science, 339, 74–78.

Janson, C.H. (1983). Adaptation of fruit morphology to dispersal agents in a neotropical forest. Science, 219, 187–188.

Janzen, D.H. (1985). On Ecological Fitting. Oikos, 45, 308.

Karger, D.N., Conrad, O., Böhner, J., Kawohl, T., Kreft, H., Soria-Auza, R.W., et al. (2017). Climatologies at high resolution for the earth’s land surface areas. Sci. Data, 4, 170–122.

Kattge, J., Bönisch, G., Díaz, S., Lavorel, S., Prentice, I.C., Leadley, P., et al. (2020). TRY plant trait database - enhanced coverage and open access. Glob. Chang. Biol., 26, 119–188.

Kissling, W.D. (2017). Has frugivory influenced the macroecology and diversification of a tropical keystone plant family? Res. Ideas Outcomes, 3, e14944.

Kissling, W.D., Balslev, H., Baker, W.J., Dransfield, J., Göldel, B., Lim, J.Y., et al. (2019). PalmTraits 1.0, a species-level functional trait database of palms worldwide. Sci. Data, 6, 178.

Lefcheck, J.S. (2016). piecewiseSEM: Piecewise structural equation modelling in r for ecology, evolution, and systematics. Methods Ecol. Evol., 7, 573–579.

Levin, S.A., Muller-Landau, H.C., Nathan, R. & Chave, J. (2003). The Ecology and Evolution of Seed Dispersal: A Theoretical Perspective. Annu. Rev. Ecol. Evol. Syst., 34, 575–604.

Lim, J.Y., Svenning, J.C., Göldel, B., Faurby, S. & Kissling, W.D. (2020). Frugivore-fruit size relationships between palms and mammals reveal past and future defaunation impacts. Nat. Commun., 11, 1–13.

Mack, A.L. (1993). The sizes of vertebrate-dispersed fruits: a neotropical-paleotropical comparison. Am. Nat., 142, 840–856.

Maruyama, P.K., Sonne, J., Vizentin-Bugoni, J., Martín González, A.M., Zanata, T.B., Abrahamczyk, S., et al. (2018). Functional diversity mediates macroecological variation in plant-hummingbird interaction networks. Glob. Ecol. Biogeogr., 27, 1186–1199.

Moles, A.T., Ackerly, D.D., Tweddle, J.C., Dickie, J.B., Smith, R., Leishman, M.R., et al. (2007). Global patterns in seed size. Glob. Ecol. Biogeogr., 16, 109–116.

Muñoz, G., Trøjelsgaard, K. & Kissling, W.D. (2019). A synthesis of animal-mediated seed dispersal of palms reveals distinct biogeographical differences in species interactions. J. Biogeogr., 46, 466–484.

Onstein, R.E., Baker, W.J., Couvreur, T.L.P., Faurby, S., Svenning, J.C. & Kissling, W.D. (2017). Frugivory-related traits promote speciation of tropical palms. Nat. Ecol. Evol., 1, 1903–1911.

Onstein, R.E., Vink, D.N., Veen, J., Barratt, C.D., Flantua, S.G.A., Wich, S.A., et al. (2020). Palm fruit colours are linked to the broad-scale distribution and diversification of primate colour vision systems. Proc. R. Soc. B Biol. Sci., 287, 20192731.

Poisot, T., Bergeron, G., Cazelles, K., Dallas, T. & Gravel, D. (2021). Global knowledge gaps in species interaction networks data. J. Biogeogr., 48, 1–17.

Running, S.W., Nemani, R.R., Heinsch, F.A., Zhao, M., Reeves, M. & Hashimoto, H. (2004). A continuous satellite-derived measure of global terrestrial primary production. Bioscience, 54, 547–560.

Sales, L.P., Kissling, W.D., Galetti, M., Naimi, B. & M Pires, M. (2021). Climate change reshapes the eco-evolutionary dynamics of a Neotropical seed dispersal system. Glob. Ecol. Biogeogr., 30, 1129–1138.

Schemske, D.W., Mittelbach, G.G., Cornell, H. V., Sobel, J.M. & Roy, K. (2009). Is there a latitudinal gradient in the importance of biotic interactions? Annu. Rev. Ecol. Evol. Syst., 40, 245–269.

Schleuning, M., Neuschulz, E.L., Albrecht, J., Bender, I.M.A., Bowler, D.E., Dehling, D.M., et al. (2020). Trait-based assessments of climate-change impacts on interacting species. Trends Ecol. Evol., 35, 319–328.

Šímová, I., Li, Y.M. & Storch, D. (2013). Relationship between species richness and productivity in plants: The role of sampling effect, heterogeneity and species pool. J. Ecol., 101, 161–170.

Sinnott-Armstrong, M.A., Donoghue, M.J. & Jetz, W.J. (2021). Dispersers and environment drive global variation in fruit colour syndromes. Ecol. Lett., 24, 1387–1399.

Sinnott-Armstrong, M.A., Downie, A.E., Federman, S., Valido, A., Jordano, P. & Donoghue, M.J. (2018). Global geographic patterns in the colours and sizes of animal-dispersed fruits. Glob. Ecol. Biogeogr., 27, 1339–1351.

Sonne, J., Vizentin-Bugoni, J., Maruyama, P.K., Araujo, A.C., Chávez-González, E., Coelho, A.G., et al. (2020). Ecological mechanisms explaining interactions within plant-hummingbird networks: Morphological matching increases towards lower latitudes. Proc. R. Soc. B Biol. Sci., 287, 20192873.

Tattersall, G.J., Arnaout, B. & Symonds, M.R.E. (2017). The evolution of the avian bill as a thermoregulatory organ. Biol. Rev., 92, 1630–1656.

Tilman, D. (1982). Resource Competition and Community Structure. (MPB-17), Volume 17. Princeton University Press.

Tobias, J.A., Ottenburghs, J. & Pigot, A.L. (2020). Avian Diversity: Speciation, Macroevolution, and Ecological Function. Annu. Rev. Ecol. Evol. Syst., 51, 533–560.

Tobias, J.A., Sheard, C., Pigot, A.L., Devenish, A.J., Yang, J. & Johnson, O. et al. (2021). AVONET: morphological, ecological and geographical data for all birds. Ecol. Lett. (this volume).

Wheelwright, N.T. (1985). Fruit size, gape width, and the diets of fruit-eating birds. Ecology, 66, 808–818.

Yap, J.Y.S., Rossetto, M., Costion, C., Crayn, D., Kooyman, R.M., Richardson, J., et al. (2018). Filters of floristic exchange: How traits and climate shape the rain forest invasion of Sahul from Sunda. J. Biogeogr., 45, 838–847.

