## Supporting information for "Global plant-frugivore trait matching is shaped by climate and biogeographic history"

**a:** Maximum trait values of each botanical country

| Response | Predictor | Estimate | DF | p-value |
|---|---|---|---|---|
| Gape size | Fruit size | 0.258 | 126 | 0.01 |
| Gape size | Palm richness | 0.466 | 128 | < 0.001 |
| Gape size | MAT | -0.14 | 128 | 0.066 |
| Fruit size | MAT | 0.26 | 127 | < 0.001 |
| Fruit size | NPP | 0.041 | 128 | 0.595 |
| Fruit size | Palm richness | 0.388 | 127 | < 0.001 |
| Palm richness | MAT | 0.368 | 129 | < 0.001 |
| Palm richness | NPP | 0.397 | 119 | < 0.001 |

**b:** Median trait values of each botanical country

| Response | Predictor | Estimate | DF | p-value |
|---|---|---|---|---|
| Gape size | Fruit size | 0.292 | 125 | 0.01 |
| Gape size | Palm richness | 0.2 | 123 | 0.005 |
| Gape size | MAT | 0.021 | 122 | 0.748 |
| Fruit size | MAT | 0.11 | 124 | 0.032 |
| Fruit size | NPP | -0.157 | 127 | 0.011 |
| Fruit size | Palm richness | -0.116 | 124 | 0.046 |
| Palm richness | MAT | 0.118 | 128 | 0.132 |
| Palm richness | NPP | 0.372 | 111 | < 0.001 |

**Table S1 |** Model coefficient (estimate) values, degrees of freedom (DF) and p-values for structural equation models in Fig. 2 and Fig. S6, which were fit with maximum (a) and median (b) botanical country values respectively.

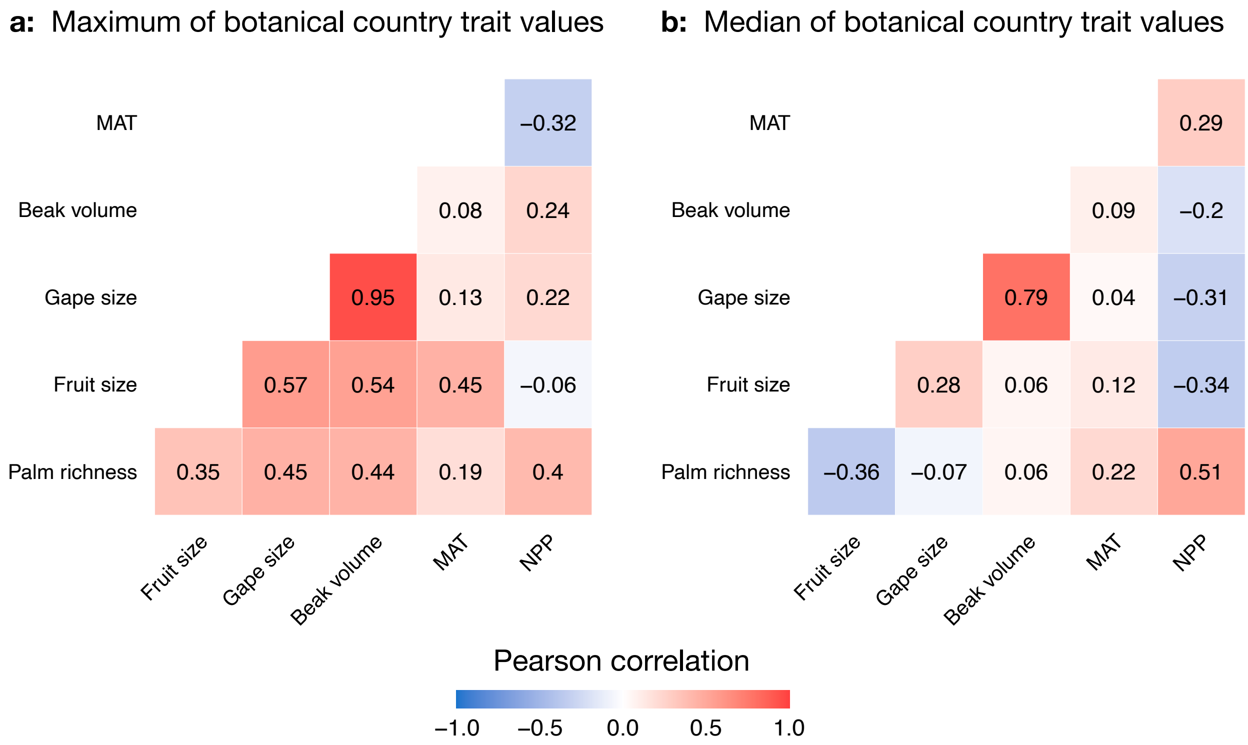


**Figure S1 |** Correlation heatmaps for log transformed bird and palm traits and climatic variables at the level of botanical countries. MAT: mean annual temperature; NPP: net primary productivity. Units: beak volume- $cm^3$; gape size- cm; fruit size- cm.

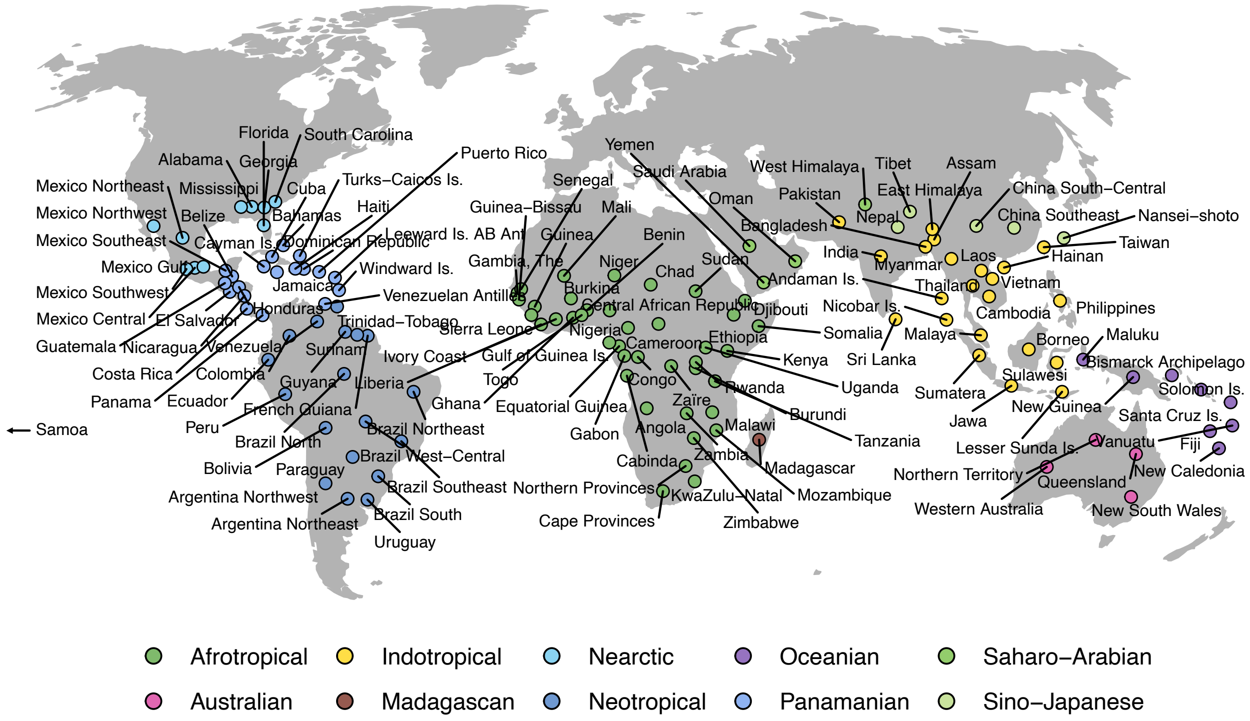


**Figure S2 |** Map of the 132 botanical countries used in the analysis (see *Methods*). Colors indicate the zoogeographic realm (*sensu* Holt *et al.* 2013) to which each botanical country belongs, dots are located at the centroids of botanical countries.

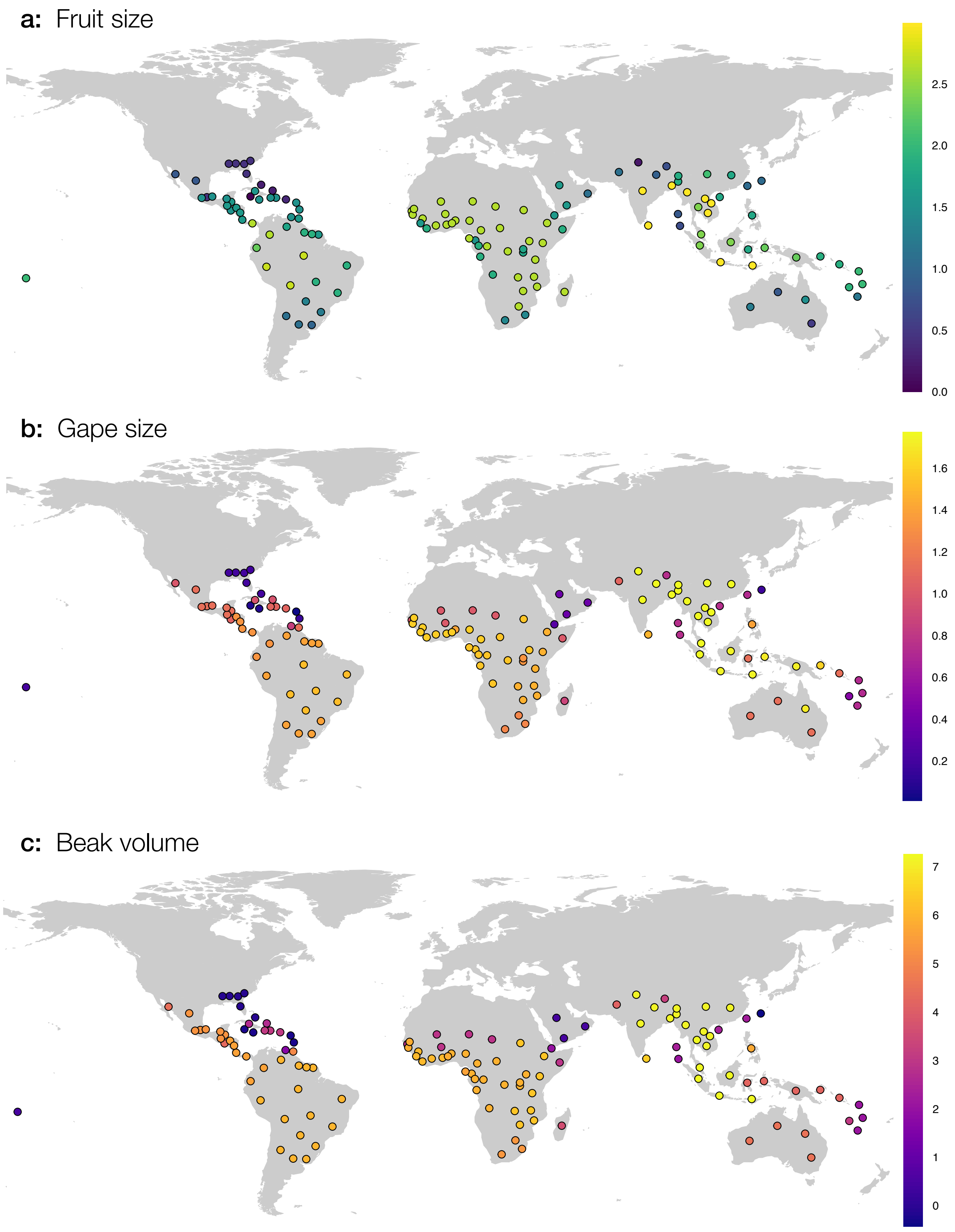


**Figure S3 |** Spatial patterns of trait variation in the *maximum* value of each botanical country. All traits were log-transformed before plotting. Units: fruit size and gape size- cm; beak volume- $cm^3$.

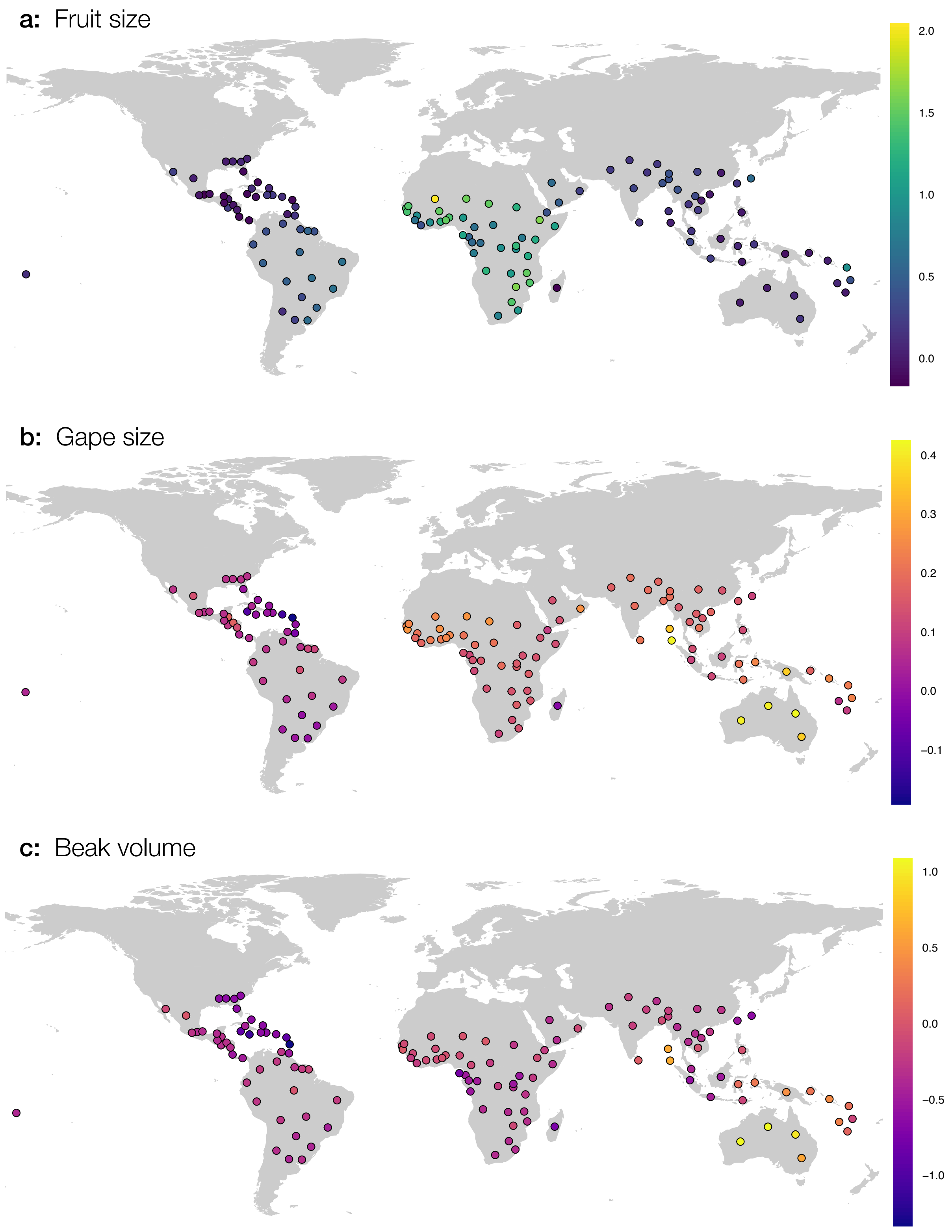


**Figure S4 |** Spatial patterns of trait variation in the *median* value of each botanical country. All traits were log-transformed before plotting. Units: fruit size and gape size- cm; beak volume- $cm^3$.

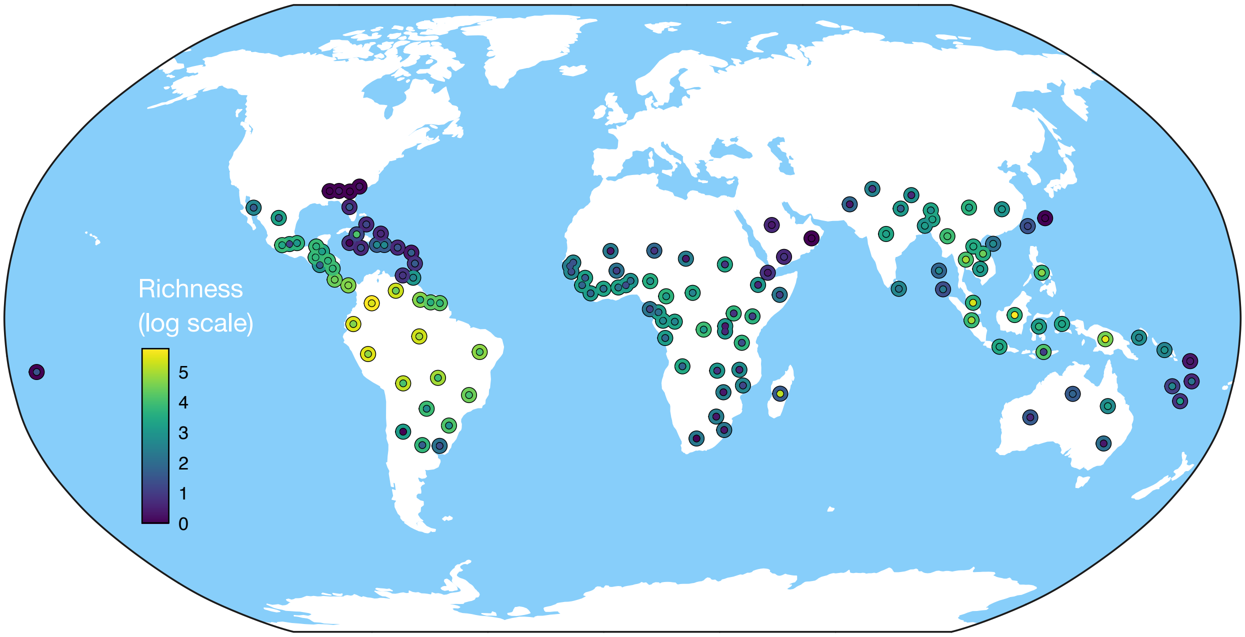


**Figure S5 |** Richness of avian frugivores and palms within each botanical country. As in Fig. 3, outer ring of points is colored by bird gape size and the inner points by fruit size. Warmer colors are higher values, note log-transformed scale. See Fig. S2 for names of each botanical country.

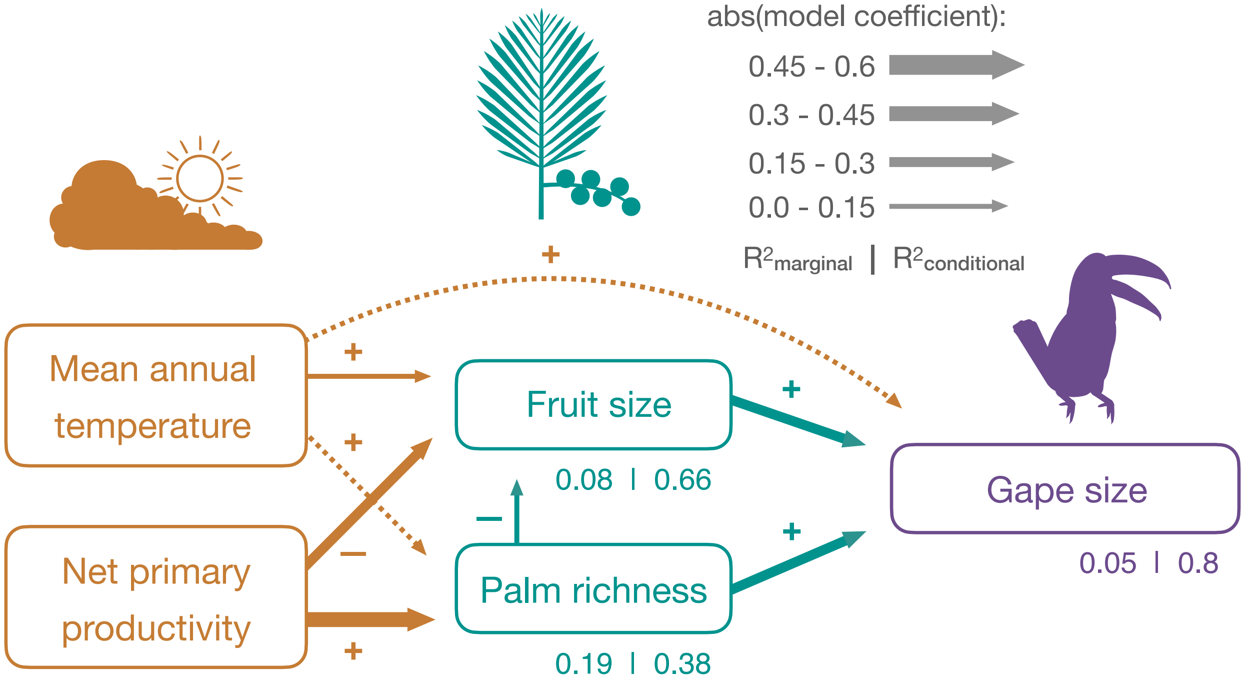


**Figure S6 |** Path diagram using median values of each botanical country (see *Methods*) in the SEM instead of maximum values (Fig. 2). Dotted paths indicate non-significant relationships, arrows are scaled to absolute values of model coefficients and signs (+/-) indicate direction of the relationship.

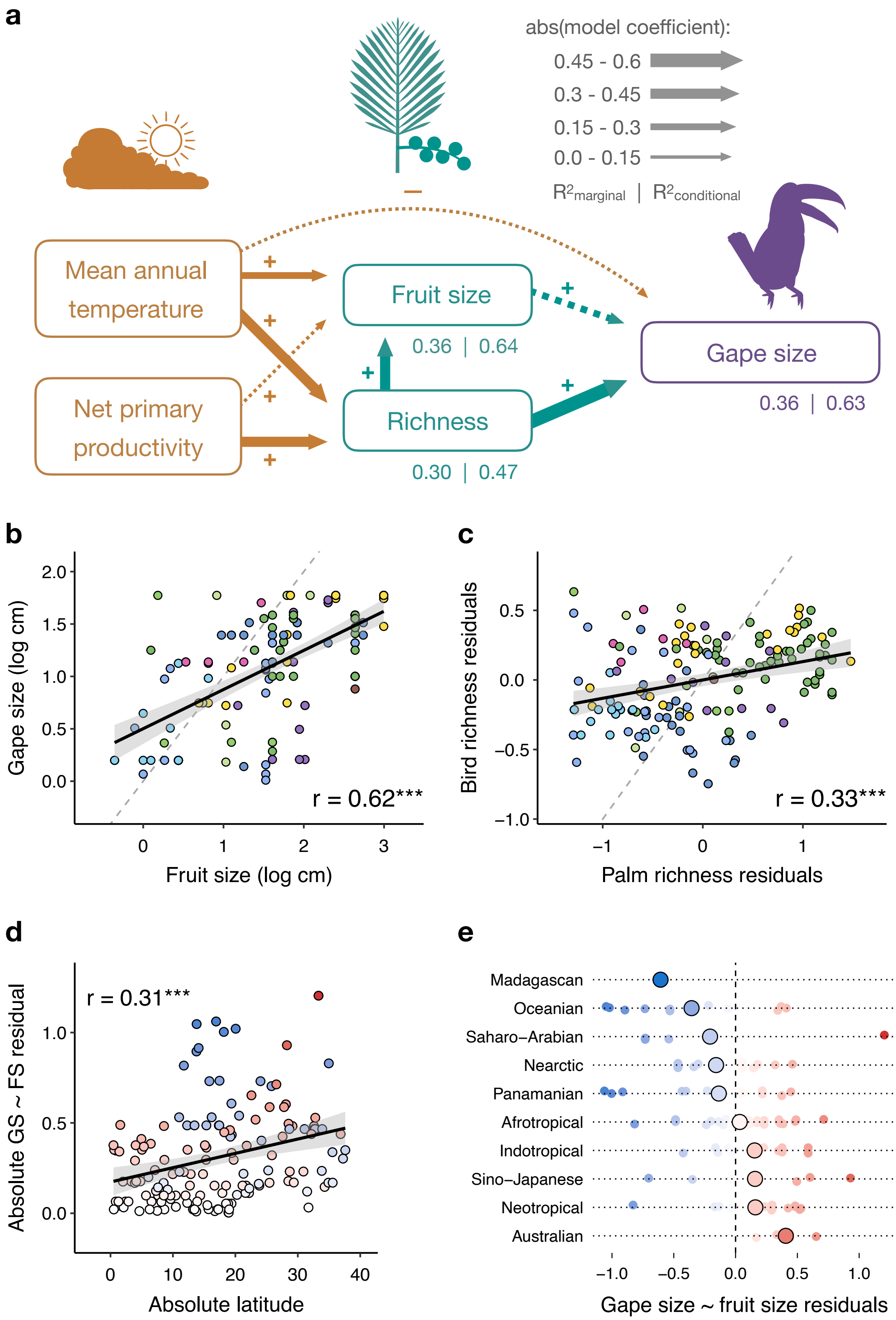


**Figure S7 |** Re-analysis of the data with the addition of 20 botanical countries with 1-2 species, using maximum trait values. Color values and line types are the same as in figures 2, 3 and 5.
